# Creative Brains Show Reduced Mid Frontal Theta

**DOI:** 10.1101/370494

**Authors:** Martijn E. Wokke, Lisa Padding, K. Richard Ridderinkhof

## Abstract

Creativity is considered to be the driving force behind innovation and progress, yet the mechanisms supporting creative thought remain elusive. In the current study, we investigated whether fluctuations in top-down control are related to creative thinking. Here, participants performed a ‘caption this’ task in which they had to provide an original and apt caption to accompany a presented picture, while EEG signals were recorded. To assess changing levels of top-down control, we made use of the strong relationship between mid frontal oscillatory activity in the theta range (4-7 HZ) and top-down control. Results demonstrate that specifically during the process of optimization and implementation of creative solutions, lower levels of mid frontal theta resulted in higher levels of creativity. In addition, increased creativity related to enhanced functional connectivity between occipital and mid frontal cortex. Together, our findings indicate that creativity benefits from a top-down induced shift towards an internally-oriented state during idea optimization and evaluation.

## Introduction

Creative thinking is often seen as the fountainhead of human progress, enabling us to go beyond existing patterns, emanating something novel and useful. Creative imagination has been considered to be the starting point of great art and science. For instance, the ideas of Nikola Tesla and the Beatles allow us to listen to Sgt. Pepper’s on the radio. In every day life, creativity also plays an important role in problem solving, adaptation to change and the discovery of new possibilities.

Although the neurocognitive mechanisms underlying creative thinking have recently attracted increasing research efforts, uncovering the neural signature of this remarkable capability has proven challenging^1,2^. Recent neuropsychological and neuroimaging findings suggest that instead of investigating where “creative neural activity” resides, it may be more fruitful to examine how neural networks interact during creative thinking^2-12^. For instance, default-mode network activity has been strongly linked to internally driven mechanisms when our focus on the external environment is diminished^13^. This internally-oriented state has been put forward to facilitate creative cognition^14-19^. Further, a considerable body of evidence links neural oscillatory mechanisms in the alpha frequency range to creative thinking^20,21^. In most studies, a higher level of alpha power in frontal, parietal, or (right-lateralized) temporal areas correlated with creative thinking^20-23^. Such effects have been interpreted as reflecting decreased externally-directed attention while supporting a more internally-oriented state during creative thinking.

Recently, the importance of top-down control networks in creative thinking has been hotly debated^24,25^. It has been shown that proper evaluation of novel ideas relies on activity in the control network^26,27^ or on the flexibility of cognitive control^28^. Other work suggests, by contrast, that reduced top-down control could in fact benefit creative processes^17,24,29^. In recent decades, mid frontal oscillatory activity in the theta range (4-7 HZ) has been associated with a broad range of top-down control processes. For instance, in the memory domain mid frontal theta has been put forward as an inhibitory mechanism biasing the competition between representations during memory retrieval^30^. In addition, mid frontal theta has been associated with the up or down regulation of activity in the default mode network^31^. In various other cognitive domains, mid frontal theta has been labeled as a robust marker of the need for top-down control^32-38^

In the present study, we investigated whether changes in mid frontal theta activity modulate processes of creative thinking. We recorded electroencephalographic (EEG) signals while participants performed a ‘caption this’ task in which they had to provide original and apt captions to accompany presented pictures. To assess the level of creativity we focused on the creative product^14,39^ and obtained creativity scores from two independent raters for each caption. We exploited the intimate relationship between mid frontal theta and top-down control by determining whether single-trial fluctuations in mid frontal theta activity and overall changes in functional connectivity (theta phase synchrony) correlated with variations in the level of creativity.

## Results

To determine the relationship between fluctuations in top-down control and creativity, we correlated per participant trial-by-trial mid frontal theta power changes (channel FCz^32^) with single trial creativity scores on a ‘caption this’ task (Figure 1)^40^ in four pre-selected time windows during the creative process. On each trial, participants provided a novel and apt one-sentence caption to accompany a presented picture (see Figure 1). Participants were instructed to press the space key the moment they came up with their solution. After the space key press the picture was removed from the screen and participants could enter the caption after a 500 ms delay period. We observed significant negative correlation coefficients between single trial theta power and single trial levels of creativity (*t*_(18)_= −3.52, *p*= 0.001, BF_−0_= 34.00, FDR-corrected *p*<0.05) exclusively in the 500 ms time period after space key press (the other three time windows −1.5 to −1 sec, −1 to −0.5 sec, and −0.5 to 0 sec relative to space key press: FDR-corrected *p*>0.05). In Figure 2a, we displayed the correlation coefficients and the t-values (between single trial theta power change and single trial levels of creativity (for illustration purposes we plotted values for all channels, while only analyzing FCz). These findings demonstrate that decreases in mid frontal theta power relates to increased creative solutions. Surprisingly, this effect was specific to the time window when the target picture was no longer visible and participants had to implement/optimize their solutions (i.e., transform their idea into an actual caption). To determine when participants finished optimizing/implementing their solutions, we computed the mean time between space key press and first entry on the keyboard. Participants took on average 3.76 seconds to start submitting their caption after space key press (SD=1.48), indicating that they were not immediately ready to submit their captions, and were most likely still optimizing (e.g., evaluating and fine-tuning) or implementing their creative solutions in this period after space key press. We further examined whether reaction times related to the level of creativity of the solutions. We therefore computed single-trial correlation coefficients between reaction time and the level of creativity for each participant. We observed no evidence for a relationship between responses and the level of creativity of the solutions (*t*_(18)_= −1.53, *p*= 0.14, BF_10_= 0.63). Next, per participant correlations coefficients between reaction time on each trial and trial-by-trial mid frontal theta power change were computed to find out whether possible differences in response preparation related to fluctuations in theta power. We observed no relation between reaction time and theta power changes (*t*_(18)_= 0.81, *p*= 0.43, BF_10_= 0.38).

**Figure 1.**
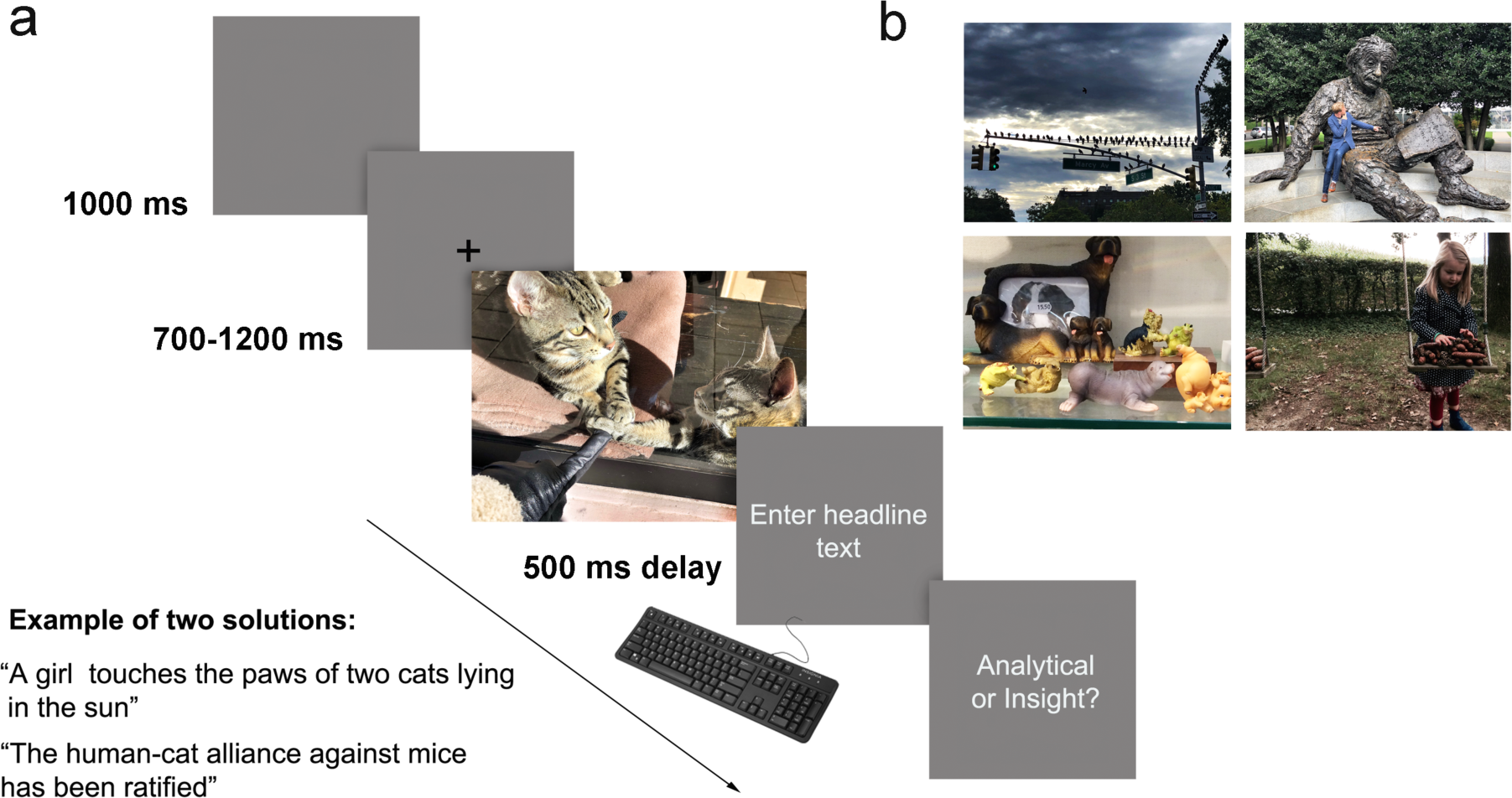
(a) After presentation of a picture participants were instructed to provide a caption to accompany the image. The image disappeared the moment participants pressed the space bar. After a short delay, participants were able to submit the caption. The trial ended probing the way participants constructed the caption. (b) Four examples of stimuli (in the experiment a total of 200 pictures were used)

**Figure 2.**
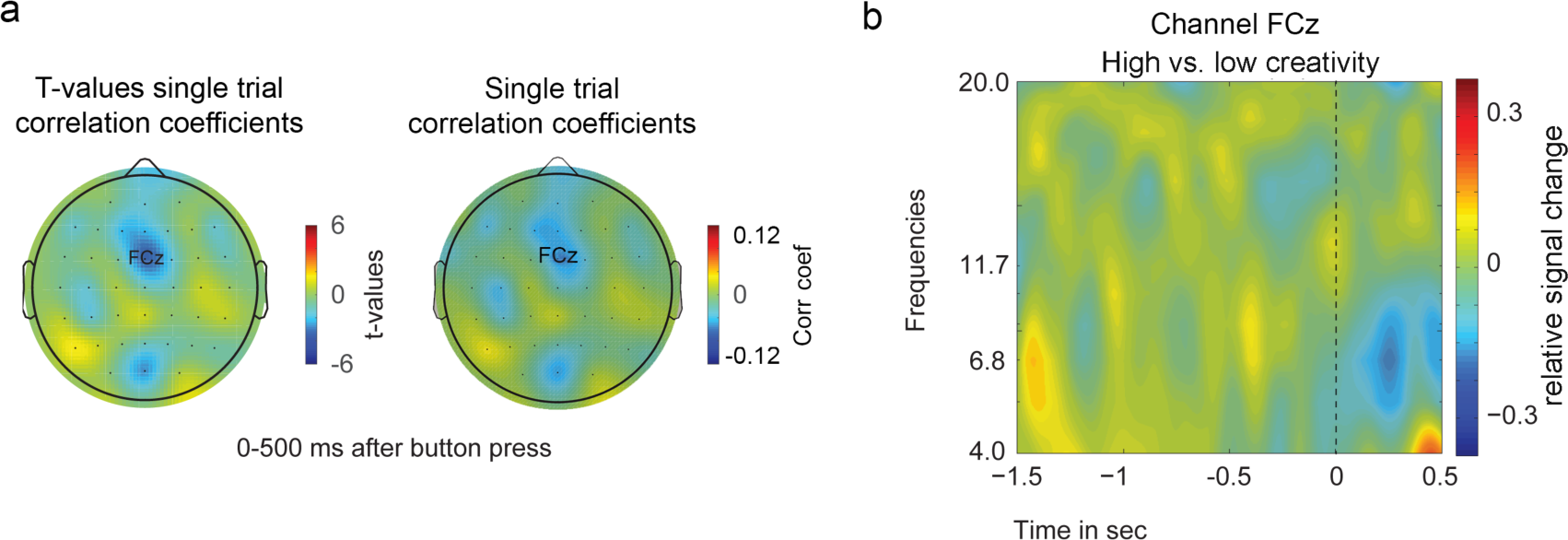
(a) We observed negative correlations between trial-by-trial change in mid frontal theta power and trial-by-trial varying levels of creativity. No correlation was found between alpha power change and levels of creativity. (b) Time-frequency plot of the difference between high (ratings >2) and low (ratings =<2) creative trials, relative to space bar press (time zero).

In light of previous findings^20-23^, we repeated the same analyses but now exploring alpha band (8-12 HZ) activity in the 500 ms after space key press. We observed no significant correlation between single trial alpha power and single trial levels of creativity in any of the time windows (FDR-corrected *p*>0.05, see Figure 3a). Because of previously observed alpha power effects in parietal, temporal and frontal electrodes we explored whether we could find a difference between high (ratings >2) and low (ratings =<2) creative trials in parietal (POz, Pz, P1, P2, P3, P4, P5, P6), temporal (TP7, TP8, T7, T8, FT7, FT8) and frontal (FCz, Fz, F1, F2, F3, F4, Fpz, Fp1, Fp2) channels (Figure 3b). No significant effects for alpha power survived the multiple comparison correction (FDR-corrected *p*>0.05).

**Figure 3.**
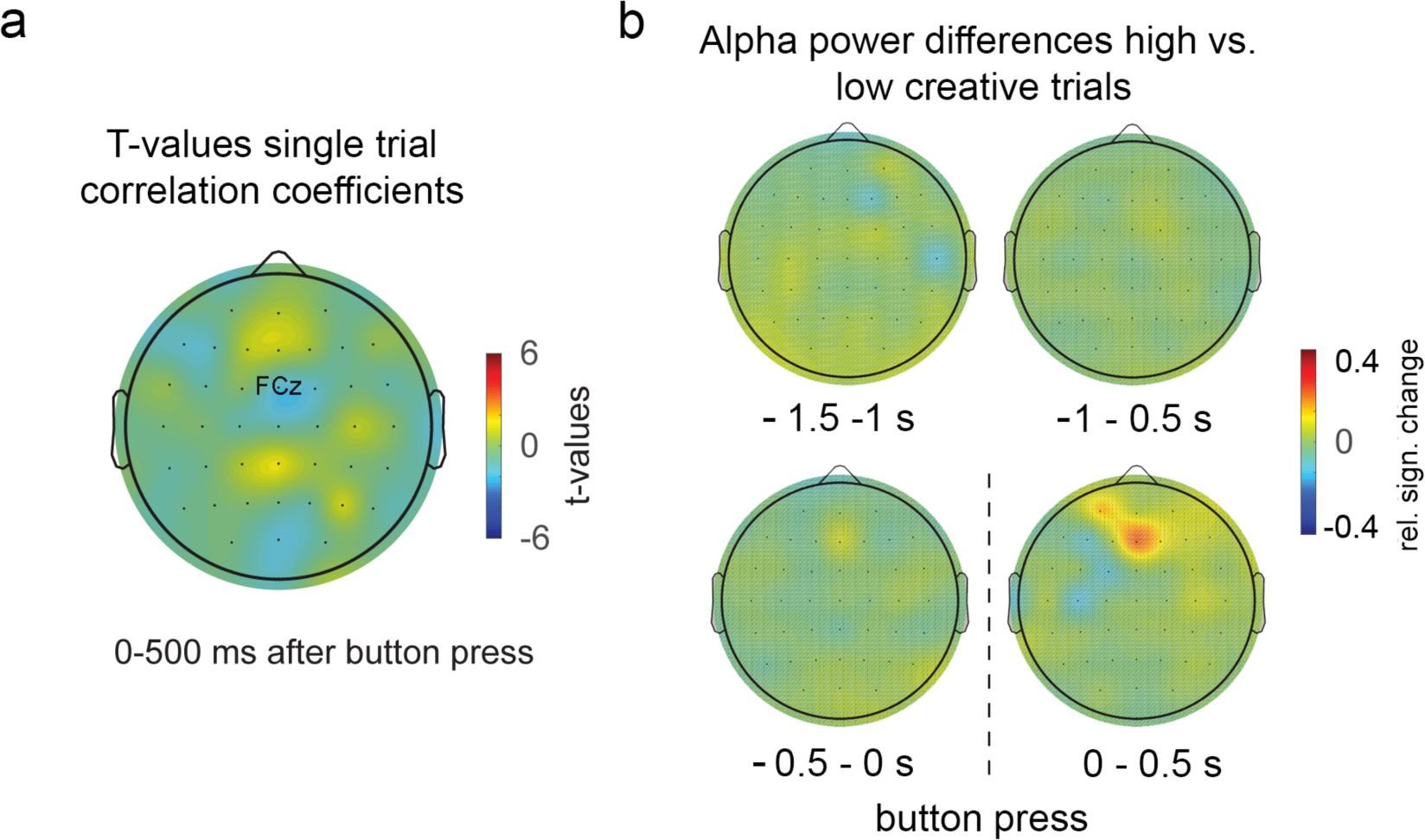
(a) We did not observe significant correlation coefficients between trial-by-trial change in alpha power and trial-by-trial varying levels of creativity. (b) Time-frequency topo plots of the difference between high and low creative trials. Despite the apparent frontal increase in alpha power for high creative trials, no result survived the correction for multiple comparisons (FDR, p >0.05).

Next, we were interested in whether changes in functional connectivity (theta phase synchrony, see methods) in the period after stimulus offset related to varying levels of creativity. We observed a significant strong positive correlation between the average level of creativity and mean changes in functional connectivity between FCz and POz (*r*= 0.60, *n*=19, *R2*= 0.36, *p* =0.007, BF_1_0= 8.58, FDR-corrected *p*<0.05). Correlations for other channels did not survive the correction for multiple comparisons (FDR>0.05), see Figure 4.

**Figure 4.**
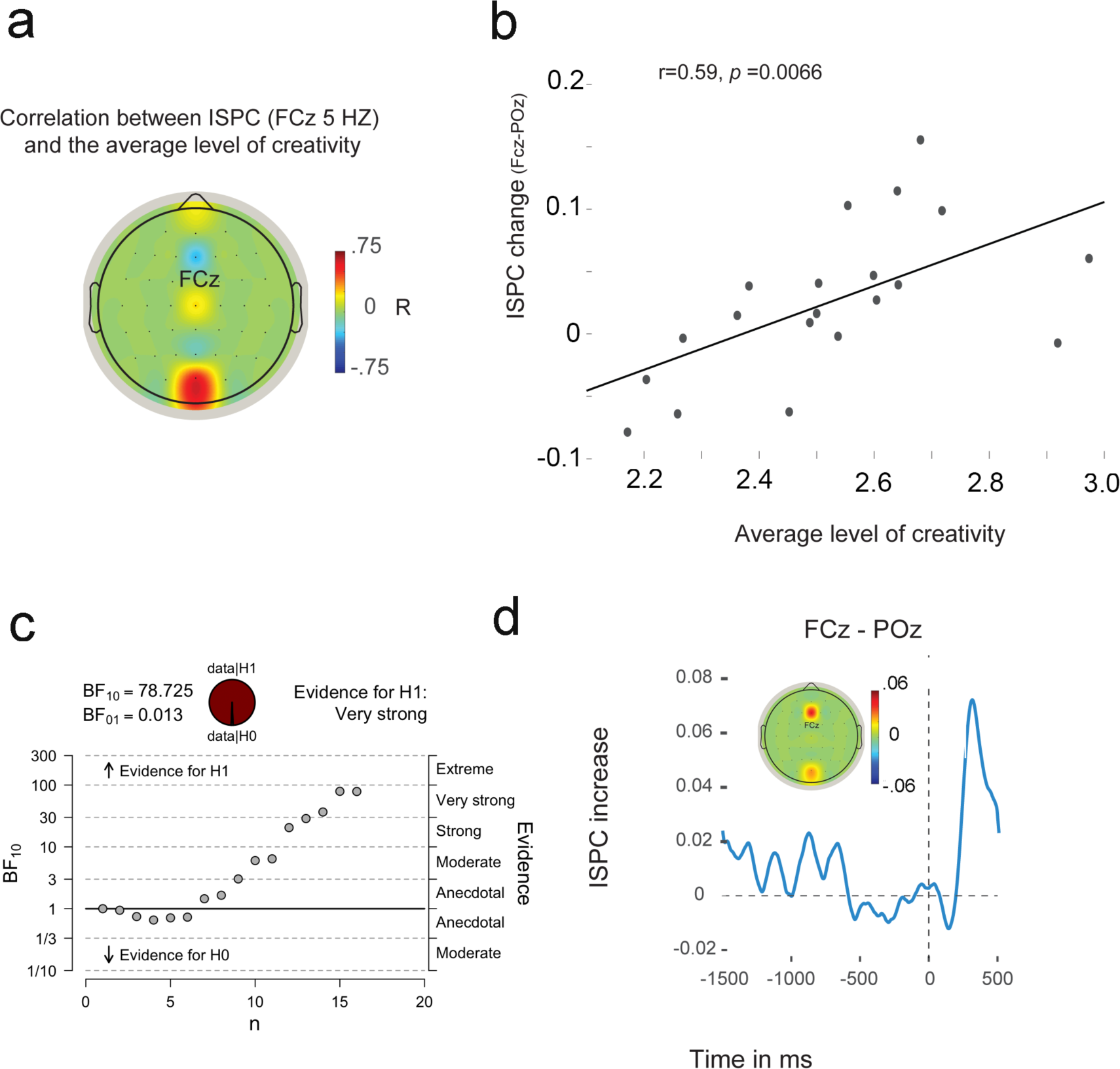
(a-b) We observed a strong positive correlation between functional connectivity (Intersite Phase Clustering [ISCP]) changes between FCz and POz and the mean level of creativity. (c) Illustration of functional connectivity change between FCz and POz over time and the topoplot of mean ISPC change between FCz and all other channels in the 500 ms after space bar press.

Finally, we explored whether the way in which solutions emerged influenced the level of creativity. We therefore asked participants to indicate whether the solution emerged through ‘sudden insight’ or through ‘analytical thought’ at the end of each trial (see methods). We observed no significant difference between ‘analytical’ (Mean 2.50, SD: 0.30) and ‘insight’ (Mean 2.61, SD: 0.25) trials with respect to average level of creativity (*t*_(18)_= 1.33, *p*= 0.198). In addition, no significant differences in alpha (t_(18)_= 0.413, *p*= 0.69) and theta power (t_(18)_= 0.662, *p*= 0.52) between approaches (insight/analytical) were found in channel FCz.

## Discussion

In the present study, we investigated whether changes in medial frontal theta oscillations related to varying levels of creativity during a ‘caption this’ task. We observed a negative relationship between trial-by-trial mid frontal theta power changes and fluctuations in creativity. In addition, we found a strong positive relationship between frontal-occipital functional connectivity change and average levels of creativity. Surprisingly, both effects were observed after participants indicated to have come up with the idea for their caption.

We have no ready explanation why these correlations were not observed during the period before space key pressing. Several factors may be at play in suppressing such correlations. For instance, an fMRI study showed that an “aha” moment of insight (in the creative process of solving anagrams) is accompanied by strong ACC activation^41^; if such aha-related effects play role in our task as well, and if such ACC activation invokes increased theta power (as might be expected from parallel literatures on error monitoring and feedback processing^42^) then this might work opposite to our anticipated ‘release-of-control’ effect. But this must remain speculative until further studies elucidate this issue.

We did find a consistent pattern of correlations in the interval between stimulus offset (space key press) and initiation of solution submission. During this relatively long period (almost 4 seconds on average) participants may engage in optimizing their creative solution. For instance, they may evaluate and fine-tune their idea further, add precision or punch, sharpen formulations to emphasize the twist, and so on. This process of further creative optimization may well benefit from reduced top-down control, thus explaining the negative correlation between creativity and mid frontal theta.

Along somewhat similar lines, an alternative explanation of the negative relationship between mid frontal theta power and the level of creativity could stem from uncertainty about the creative quality of their solution. To the extent that participants are sensitive to the level of creativity of their own solutions, trial-by-trial fluctuations of certainty about the solution may result, in turn inducing trial-by-trial fluctuations in top-down control^32^. If this would be the case, less creative solutions would be accompanied by enhanced mid frontal theta, thereby rendering the observed theta effect somewhat epiphenomenal to the actual creative process. Effects of uncertainty about the solution could become apparent in the period after space key press, when the idea needs to be phrased and evaluated. Note that as uncertainty about the solution increases, reaction times typically also increase^43^. However, here we did not observe a relationship between creativity and the time it took participants to start submitting their solution, nor did we find a relation between reaction times and mid frontal theta changes. It therefore seems unlikely that our observed effects are related to participants’ uncertainty about the creative quality of their ideas. Complementary to the solution optimization account, the observed effects might relate to certain other processes associated less directly with the implementation of the generated ideas. Following space key press, after ideas has been generated and selected, participants needed to transform their ideas into concrete phrases. During this period, the stimulus was removed from the screen, prompting attention to become internally oriented. It has been demonstrated that occipital-frontal functional connectivity measured in the theta range was associated with keeping (perceptual) information “in mind”^44-46^. Von Stein and Sarnthein^47^ observed a relationship between increased long-range (frontal-posterior) theta synchronization and working-memory retention. In their study, EEG signals were analyzed from the part of the experiment when no external stimulus was present, allowing them to observe activity during entirely internal processing. Similarly to our study, in this phase of a trial internal activity was still partly controlled by the previously presented stimulus. Increased theta synchronization between frontal and posterior cortex were specifically found during the period after stimulus offset, when perceptual information needed to be kept active or imagined.

In tune with this account, another line of work demonstrates that default mode network (DMN) activity, strongly associated with an internally-oriented state, negatively correlates with mid frontal theta activity^31,48,49^. In both resting state and task active settings, concurrent EEG/fMRI studies showed that reduced DMN activity related to increased mid frontal theta activity. Although the exact relationship between DMN activity and mid frontal theta remains to be further investigated, it is likely that DMN deactivation could be an indicator of theta-related processes^50^. In the present study, creativity in inventing the caption might benefit from stronger shifts towards internally-oriented operations (increased occipital-frontal functional connectivity, and decreased mid frontal theta) during implementation of the solution,.

Neural oscillations have been put forward as a mechanism that supports information integration and communication between large-scale neural networks^51-55^. Creativity has recently been associated with large-scale network interactions^7,12^, and the integration of distributed neural representation^56^. Recently, it has been hypothesized that alpha oscillations are pivotal in the gating of the flow of information via inhibition of task-irrelevant network activity^57-60^. For instance, studies showed that a shift of attention to either the left or right visual hemifield decreases alpha in the contralateral hemisphere, and depending on task demands, alpha power increased in dorsal stream when a task relied on ventral stream processing^60,61^. Previous studies demonstrated that activity in the alpha band related to the divergent-convergent dichotomy^62^.. Interestingly, increased alpha activity has been associated with an increase in internally-oriented attention, suggesting the necessity of active inhibition of external sensory input for internally driven mental operations^63,63^. However, differences in alpha power have also been strongly linked to the amount of effort or attention that is allocated to the task at hand^60^. Indeed, some findings indicate that the difference between divergent and convergent thinking stems from a difference in task demands^46,65^. In the present study, we focused our analyses on mid frontal theta activity, however, when examining alpha power we did not observe a relation between creativity and alpha power fluctuations (see Figure 3). Similarly, Boot et al.^66^ did not observe a relation between convergent-divergent thinking and alpha activity. The authors argued that the use of an event-related design ‘canceled out the effect of differences in the demands that (the) different tasks place on cognitive resources, rather than the fast-acting processes underlying the creative processes’. As in the present study, Boot et al.^66^ used an event related design that investigated trial-by-trial fluctuations during a task that probed divergent vs. convergent thinking (thereby keeping processing demands constant). Interestingly, Boot et al. observed a selective decrease of delta band activity associated with divergent thinking. Previously, decreased delta-band activity over fronto-central regions has been linked to internally oriented processes and default mode network activity^9,67^. These findings are in line with the present results indicating the importance of internally-oriented network activity and creativity.

One of the major problems when attempting to uncover the neural basis of creativity is to adequately capture and isolate creative thinking. Most psychometric tests currently used are based on dichotomies between for instance divergent-convergent thinking and defocused-focused attention^14,68,69,70^. However, creative thinking has been associated with both sides of such divisions, making it challenging to separate instances of creative thinking from ‘normal thinking’. The main problem is to determine the criterion distinguishing creative vs. normal thinking. Some have argued that research into creativity should therefore focus on the ‘creative product’ itself^39^. In the present study, we focused on the creative solution without looking into traditional convergent-divergent or defocused-focused attention dichotomies. This approach has clear limitations, for instance, by creating a more fuzzy/subjective definition of creativity. However, we believe that this approach is more ecologically valid. As depicted above, dichotomies such as divergent-convergent thinking or defocused-focused attention might result in clear divisions on one scale, however, when trying to capture and isolate creative thinking and distinguish it from ‘normal thinking’ such divisions could result in throwing away the baby with the bathwater. When using a creative product approach, it is critical that a large variation of tasks, groups and methods are being used, while keeping possible contaminating factors (such as task load) constant. Therefore, it would be interesting and important to investigate whether the present results generalize to different tasks and settings that capture different aspects of creative thinking.

In sum, the present findings demonstrate that lower levels of frontal theta related to creative performance. In addition, we observed that increased creativity was associated with enhanced long-range functional connectivity between occipital and mid frontal cortex. These results contribute to a growing amount of evidence linking creativity to large-scale network dynamics, orchestrating the balance between internally and externally oriented network activity. These findings could be important in (the) light of ongoing technological developments (i.e., portable electronic devices) reducing internal reflection by biasing external attention^71^.

## Materials and Methods

### Participants

Twenty-four participants (mean age= 21.5, SD= 3.2) took part in this study for financial compensations. All participants had normal or corrected-to-normal vision, and all were naïve to the purpose of the experiment. Four participants were excluded because of excessive noise in more than half of the trials (due to our long epochs, see below). One participant was excluded for not performing the task correctly. A total of nineteen participants were included for all further analyses. All procedures complied with international laws and institutional guidelines and were approved by the Ethics Committee of the Psychology department of the University of Amsterdam, and all participants provided their written informed consent prior to the experiment.

### Task design

Pictures were presented full screen (1024*768 pixels) on a 17-inch DELL TFT monitor with a refresh rate of 60 Hz. The monitor was placed at a distance of ∼90 cm in front of each participant so that one centimeter subtended a visual angle of 0.64°. Each trial started with a blank (1000 ms) after which a fixation-cross appeared (jittered between 700-1200 ms, in steps of 100 ms). Next, a picture was presented, which consisted of a scene or an event (see Figure 1). On each trial, we instructed participants to provide an apt and original one-sentence caption to accompany the presented picture. Participants were instructed to press the space key on a keyboard at the moment they came up with the content of their caption. After the space key press, the picture disappeared and participants could type in their response after a 500 ms delay period. Each caption was submitted by pressing the enter key. The trial ended with a question about how the caption emerged. Participants could respond by pressing the ‘1’ key in case the caption was thought up of in an analytical manner or the ‘2’ key in case the caption emerged through sudden insight. Sudden insight was described as a caption that ‘*popped into mind’*, while analytic solutions were described as coming into being after ‘*analytical thought and reasoning*’.

The experiment lasted ∼2 hours and consisted of 200 trials divided into 10 blocks. The 200 pictures (selected during a pilot experiment) were presented in pseudo-random order. Stimuli were presented using Presentation (Neurobehavioral Systems).

At the end of the experiment, two naïve participants rated the level of creativity of each caption (3800 trials in total) by awarding a score between 1-4, where 1 represented a very low level of creativity and 4 an extremely high level of creativity. The raters were instructed that a creative caption was ‘*a caption that was clearly connected to the image, while describing the picture in an original and novel way. Further, the content of the caption should go beyond the information provided by the image’*. Low creativity was described as ‘*providing a purely factual depiction of the presented image’* and/or *‘clearly not connected to the picture at all*’. We calculated inter-rater reliability (Cohen’s kappa), and we observed a fair amount^72^ of agreement (κ = .267, p < .0005) between the two raters. Next, we excluded all trials on which the raters differed >2 points from further analyses (a total of 15 trials across all participants).

### EEG measurements and analyses

We recorded and sampled EEG signals at 1048 Hz using a Biosemi ActiveTwo 64-channel system, with four additional electrodes for horizontal and vertical eye-movements, each referenced to their counterpart (Biosemi – Amsterdam, The Netherlands). High-pass filtering (0.5 HZ), low-pass filtering (100 HZ) and a notch filter (50 HZ) were used. Next, eye movements were corrected on the basis of Independent Component Analysis^73^, after which the signal was down-sampled to 512 Hz. We epoched the data −3 to + 1 sec surrounding space key response and removed trials containing irregularities due to EMG or other artifacts by visually inspecting all trials. To increase spatial specificity and to filter out deep sources we converted the data to spline Laplacian signals^40^. Per participant and per electrode we subtracted the average of all trials from each individual trial to obtain the non-phase-locked power^74,75,76^. Next we used a sliding window Fourier transform^77^, window length: 400 ms, step size: 50 ms, to calculate the time-frequency representations of the EEG power (spectrograms) for each channel and each trial. We used a single Hanning taper for the frequency range 4–25 Hz (bin size: 1 Hz^77^). Power modulations were characterized as the percentage of power change at a given time and frequency bin relative to baseline power value for that frequency bin. The baseline was calculated as the mean power in the pre-stimulus interval (from −0.3 to 0 s relative to picture onset). All signal processing steps were performed using Brain Vision Analyzer (BrainProducts), Matlab (Matlab 12.1, The MathWorks Inc.), **X** ^code55^ and Fieldtrip^78^.

It has been proposed that creativity relies on binding of previously unconnected representations, established by patterns of neural activity^56^ Therefore, creativity might require long-range coordination between distant cortical regions^76^. To investigate measures of interregional functional connectivity, we assessed the consistency of the difference of time–frequency phase values between two channels in the theta band across trials (Intersite Phase Clustering (ISPC)^55^). We used FCz as our ‘seed’ electrode and paired that electrode with central electrodes Oz, POz, Cz, Fz, AFz Fpz, and dorsolateral electrodes F4/F3. We used the same preprocessing steps as described above for the time-frequency analyses. We used a baseline period of −300-0 ms before picture onset.

In this experiment, we were specifically interested in the relationship between fluctuations in mid frontal (FCz) theta power, changes in theta phase synchrony and varying levels of creativity. We therefore correlated single trial mean theta power changes (compared to baseline) in a time window of −1.5 to −1 sec, −1 to −0.5 sec, −0.5 to 0 sec, and 0 to 0.5 sec relative to space key press with single trial creativity scores per participant (time windows were based on space key press times from a pilot study). Next, we tested whether correlation coefficients differed from zero using t-tests^79^. We pre-selected these four time windows to reduce the amount of comparisons in our analyses, while maintaining the ability to distinguish between different phases of the task (no data before −1.5 sec prior to space key press was selected because the task was self paced, thereby creating unequal trial lengths). We tested these correlations against zero using one-sample t-tests. Next, we correlated mean levels of creativity with theta phase synchrony values (using FCz as a ‘seed’ electrode, see above), using our theta power results to base our time widow of interest upon (to limit the amount of comparisons). Because we examined the consistency of the difference of time–frequency phase values between two channels (i.e., here phase differences between two channels on a single trial are uninformative, as we are interested in the clustering of this difference), we therefore correlated ISPC changes (compared to baseline) with mean levels of creativity (across trials). We corrected for multiple comparisons by adjusting the p value by fixing the false discovery rate (FDR) at 0.05 ^80^.

Finally, we tested whether insight vs. analytical solutions differed in the level of creativity. We tested mean levels of creativity for trials that were judged as analytical versus trials that were judged as insight trials, using a paired t-test (two-sided). Further, we determined whether theta power differed between insight vs. analytical solutions using a paired t-test (two-sided).

All individuals identifiable in the images used in figure 1 provided informed consent for publication in an online open-access publication, and in the case of individuals under the age of 18 years, informed consent has been obtained from a parent and legal guardian.

## Acknowlegments

We would like to thank Djoni de Vos for helping with the stimuli selection and piloting the experiment. This work was supported by the Freek and Hella de Jonge Creative Mind Prize (MEW), the Amsterdam Brain and Cognition Talent Grant (MEW) and the European Union’s Horizon 2020 research and innovation programme under the Marie Skłodowska-Curie grant agreement (Meta_Mind -DLV-704361 to MEW).

## Author Contributions

M.E.W. designed the research; L.P. performed the research; M.E.W., K.R.R. and L.P. analyzed the data; M.E.W. and K.R.R. wrote the manuscript.

## Competing Interests

The authors declare no competing interests.

## Notes

#### Summary of Updates

Received reviews substantially improved the manuscript.

## References

1. Dietrich, A., & Kanso, R. A review of EEG, ERP and neuroimaging studies of creativity and insight. Psychological Bulletin, 136, 822–848 (2010).

2. Dietrich, A., & Haider, H. A Neurocognitive Framework for Human Creative Thought. Frontiers in Psychology, 7, 1–7 (2017).

3. Beaty, R. E., Benedek, M., Wilkins, R. W., Jauk, E., Fink, A., Silvia, P. J., Hodges, D. A., Koschutnig, K., & Neubauer, A. C. Creativity and the default network: A functional connectivity analysis of the creative brain at rest. Neuropsychologia, 64, 92–98 (2014).

4. Jung, R. E., Grazioplene, R., Caprihan, A., Chavez, R. S., & Haier, R. J. White matter integrity, creativity, and psychopathology: disentangling constructs with diffusion tensor imaging. PLoS One, 5(3), 9818 (2010).

5. Chrysikou, E. G., Novick, J. M., Trueswell, J. C., & Thompson-Schill, S. L. The other side of cognitive control: Can a lack of cognitive control benefit language and cognition? Topics in Cognitive Science, 3, 253–256 (2011).

6. Mell, J. C., Howard, S. M., & Miller, B. L. Art and the brain: The influence of frontotemporal dementia on an accomplished artist. Neurology, 60, 1707–1710 (2003).

7. Zabelina, D. L., & Andrews-Hanna, J. R. Dynamic network interactions supporting internally-oriented cognition. Current opinion in neurobiology, 40, 86–93 (2016).

8. McMillan, R., Kaufman, S. B., & Singer, J. L. Ode to positive constructive daydreaming. Frontiers in psychology, 4, 626 (2013).

9. Baird, B., Smallwood, J., Mrazek, M. D., Kam, J. W., Franklin, M. S., & Schooler, J. W. Inspired by distraction: mind wandering facilitates creative incubation. Psychological science, 23(10), 1117–1122 (2012).

10. Mok, L. W. The interplay between spontaneous and controlled processing in creative cognition. Frontiers in human neuroscience, 8, 663 (2014).

11. Zedelius, C. M., & Schooler, J. W. The richness of inner experience: relating styles of daydreaming to creative processes. Frontiers in psychology, 6, 2063 (2016).

12. Beaty, R. E., Kenett, Y. N., Christensen, A. P., Rosenberg, M. D., Benedek, M., Chen, Q., Fink, A. Qiu, J., Kwapil, T. R., Kane, M. J., & Silvia, P. J. Robust prediction of individual creative ability from brain functional connectivity. Proc of the Nat Ac of Sci, 115(5), 1087–1092 (2018).

13. Buckner, R. L., Andrews-Hanna, J. R., & Schacter, D. L. The brain’s default network. Annals of the New York Academy of Sciences, 1124(1), 1–38 (2008).

14. Dietrich, A. Who’s afraid of a cognitive neuroscience of creativity? Methods, 42(1), 22–27 (2007).

15. Kaufman, S. B., DeYoung, C. G., Gray, J. R., Jiménez, L., Brown, J., & Mackintosh, N. Implicit learning as an ability. Cognition, 116(3), 321–340 (2010).

16. Carson, S. H. Cognitive disinhibition, creativity, and psychopathology. The Wiley handbook of genius, 198–221 (2014).

17. Chrysikou, E. G., Hamilton, R. H., Coslett, H. B., Datta, A., Bikson, & M., Thompson-Schill, S. L. Noninvasive Transcranial Direct Current Stimulation Over the Left Prefrontal Cortex Facilitates Cognitive Flexibility in Tool Use. Cognitive Neuroscience. 4(2), 81–89 (2013).

18. Fox, M. D., Snyder, A. Z., Vincent, J. L., Corbetta, M., van Essen, D. C., & Raichle, M. E. The human brain is intrinsically organized into dynamic, anticorrelated functional networks. Proc of the Nat Ac of Sci, 102(27), 9673–9678 (2005).

19. Wokke, M. E., Talsma, L. J., & Vissers, M. E. Biasing neural network dynamics using noninvasive brain stimulation. Frontiers in systems neuroscience, 8, 246 (2015).

20. Fink, A., & Benedek, M. EEG alpha power and creative ideation. Neuroscience & Biobehavioral Reviews, 44, 111–123 (2014).

21. Benedek, M., Bergner, S., Könen, T., Fink, A., & Neubauer, A. C. EEG alpha synchronization is related to top-down processing inconvergent and divergent thinking. Neuropsychologia, 49(12), 3505–3511 (2011).

22. Lustenberger, C., Boyle, M. R., Foulser, A. A., Mellin, J. M., & Fröhlich, F. Functional role of frontal alpha oscillations in creativity. Cortex, 67, 74–82 (2015)

23. Luft, C. D. B., Zioga, I., Thompson, N. M., Banissy, M. J., & Bhattacharya, J. Right temporal alpha oscillations as a neural mechanism for inhibiting obvious associations. Proceedings of the National Academy of Sciences, 115(52), (2018)

24. Amer, Tarek, Karen L. Campbell, and Lynn Hasher. “Cognitive control as a double-edged sword.” Trends in cognitive sciences 20.12, 905–915 (2016)

25. Beaty, Roger E., et al. “Creative cognition and brain network dynamics.” Trends in cognitive sciences 20.2, 87–95 (2016).

26. Ellamil, Melissa, et al. “Evaluative and generative modes of thought during the creative process.” Neuroimage 59.2 1783–1794, (2012).

27. Beaty, R. E., Benedek, M., Kaufman, S. B., & Silvia, P. J. Default and executive network coupling supports creative idea production. Scientific reports, 5, 10964 (2015).

28. Zabelina, Darya L., and Michael D. Robinson. “Creativity as flexible cognitive control.” Psychology of Aesthetics, Creativity, and the Arts 4.3 136 (2010).

29. Chrysikou, Evangelia G., and Sharon L. Thompson-Schill. “Dissociable brain states linked to common and creative object use.” Human brain mapping 32.4, 665–675 (2011).

30. Hsieh, L. T., & Ranganath, C. Frontal midline theta oscillations during working memory maintenance and episodic encoding and retrieval. Neuroimage, 85, 721–729 (2014).

31. Scheeringa, R., Bastiaansen, M. C., Petersson, K. M., Oostenveld, R., Norris, D. G., & Hagoort, P. Frontal theta EEG activity correlates negatively with the default mode network in resting state. International journal of psychophysiology, 67(3), 242–251 (2008).

32. Cavanagh, J. F., & Frank, M. J. Frontal theta as a mechanism for cognitive control. Trends in Cognitive sciences, 18, 8 (2014).

33. Van de Vijver, I., Ridderinkhof, K. R., & Cohen, M. X. Frontal oscillatory dynamics predict feedback learning and action adjustment. Journal of Cognitive Neuroscience, 23(12), 4106–4121 (2011).

34. Van Driel, J., Swart, J. C., Egner, T., Ridderinkhof, K. R., & Cohen, M. X. (No) time for control: frontal theta dynamics reveal the cost of temporally guided conflict anticipation. Cognitive, Affective, & Behavioral Neuroscience, 15(4), 787–807 (2015).

35. Razoumnikova, O. M. Creativity related cortex activity in the remote associates task. Brain Research Bulletin, 63(1-3), 96–102 (2007).

36. Sauseng, P., Klimesch, W., Freunberger, R., Pecherstorfer, T., Hanslmayr, S., & Doppelmayr, M. Relevance of EEG alpha and theta oscillations during task switching. Experimental Brain Research, 170(3), 295–301 (2006).

37. Cohen, M. X., & Cavanagh, J. F. Single-trial regression elucidates the role of prefrontal theta oscillations in response conflict. Frontiers in Psychology, 2, 30 (2011).

38. Wokke, M. E., Cleeremans, A., & Ridderinkhof, K. R. Sure I’m sure: Prefrontal oscillations support metacognitive monitoring of decision-making. Journal of Neurosci, 37(4), 781–789 (2017).

39. MacKinnon, D. W. Some critical issues for future research in creativity. Frontiers of creativity research: Beyond the basics, 120–130 (1987).

40. Nusbaum, E. C., Silvia, P. J., & Beaty, R. E. (2017). Ha ha? Assessing individual differences in humor production ability. Psychology of Aesthetics, Creativity, and the Arts, 11(2), 231.

41. Aziz-Zadeh, L., Kaplan, J. T. and Iacoboni, M. (2009), “Aha!”: The neural correlates of verbal insight solutions. Hum. Brain Mapp., 30: 908–916.

42. Ridderinkhof, K. R., Ullsperger, M., Crone, E. A., & Nieuwenhuis, S. (2004). The role of medial frontal cortex in cognitive control. Science, 306, 443–447

43. Weidemann, C. T., & Kahana, M. J. (2016). Assessing recognition memory using confidence ratings and response times. Royal Society open science, 3(4), 150670.

44. Andrews-Hanna, J. R., Smallwood, J., & Spreng, R. N. The default network and self-generated thought: component processes, dynamic control, and clinical relevance. Annals of the New York Academy of Sciences, 1316(1), 29–52 (2014).

45. Sarnthein, J., Petsche, H., Rappelsberger, P., Shaw, G. L., & Von Stein, A. Synchronization between prefrontal and posterior association cortex during human working memory. Proc of the Nat Ac of Sci, 95(12), 7092–7096 (1998).

46. Chou, W. C., Duann, J. R., She, H. C., Huang, L. Y., & Jung, T. P. Explore the functional connectivity between brain regions during a chemistry working memory task. PloS one, 10(6), e0129019 (2015).

47. Von Stein, A., & Sarnthein, J. Different frequencies for different scales of cortical integration: from local gamma to long range alpha/theta synchronization. International journal of psychophysiology, 38(3), 301–313 (2000).

48. White, T. P., Jansen, M., Doege, K., Mullinger, K. J., Park, S. B., Liddle, E. B., … & Liddle, P. F. (2013). Theta power during encoding predicts subsequent-memory performance and default mode network deactivation. Human brain mapping, 34(11), 2929–2943.

49. Scheeringa, R., Petersson, K. M., Oostenveld, R., Norris, D. G., Hagoort, P., & Bastiaansen, M. C. (2009). Trial-by-trial coupling between EEG and BOLD identifies networks related to alpha and theta EEG power increases during working memory maintenance. Neuroimage, 44(3), 1224–1238.

50. Hsieh, L. T., & Ranganath, C. (2014). Frontal midline theta oscillations during working memory maintenance and episodic encoding and retrieval. Neuroimage, 85, 721–729.

51. Buzsaki, G. (2006). Rhythms of the Brain. Oxford University Press.

52. Fries, P. A mechanism for cognitive dynamics: neuronal communication through neuronal coherence. Trends in cognitive sciences, 9(10), 474–480 (2005).

53. Hipp, J. F., Engel, A. K., & Siegel, M. Oscillatory synchronization in large-scale cortical networks predicts perception. Neuron, 69(2), 387–396 (2011).

54. Siegel, M., Donner, T. H., & Engel, A. K. Spectral fingerprints of large-scale neuronal interactions. Nature Reviews Neuroscience, 13(2), 121 (2012).

55. Cohen, M. X. (2014). Analyzing neural time series data: theory and practice. MIT press.

56. Thagard, P., & Stewart, T. C. The AHA! Experience: Creativity Through Emergent Binding in Neural Networks. Cognitive Science, 35, 1–33 (2010).

57. Mathewson, K. E., Beck, D. M., Ro, T., Maclin, E. L., Low, K. A., Fabiani, M., & Gratton, G. Dynamics of alpha control: preparatory suppression of posterior alpha oscillations by frontal modulators revealed with combined EEG and event related optical signal. Journal of cognitive neuroscience, 26(10), 2400–2415 (2014).

58. Jensen, O. & Mazaheri, A. “Shaping functional architecture by oscillatory alpha activity: gating by inhibition.” Frontiers in human neuroscience, 4, 186 (2010).

59. Klimesch, W. EEG alpha and theta oscillations reflect cognitive and memory performance: a review and analysis. Brain research reviews, 29(2-3), 169–195 (1999).

60. Jokisch, D., & Jensen, O. Modulation of gamma and alpha activity during a working memory task engaging the dorsal or ventral stream. Journal of Neuroscience, 27(12), 3244–3251 (2007).

61. Wokke, M. E., Scholte, H. S., & Lamme, V. A. Opposing dorsal/ventral stream dynamics during figure-ground segregation. Journal of cognitive neuroscience, 26(2), 365–379 (2014).

62. Jauk, E., Benedek, M., & Neubauer, A. C. (2012). Tackling creativity at its roots: Evidence for different patterns of EEG alpha activity related to convergent and divergent modes of task processing. International Journal of Psychophysiology, 84(2), 219–225 (2012).

63. Cooper, N. R., Croft, R. J., Dominey, S. J., Burgess, A. P., & Gruzelier, J. H. (2003). Paradox lost? Exploring the role of alpha oscillations during externally vs. internally directed attention and the implications for idling and inhibition hypotheses. International Journal of Psychophysiology, 47(1), 65–74.

64. Wokke, M. E., & Ro, T. (2018). Anterior prefrontal cortex mediates implicit inferences. bioRxiv, 390427.

65. Zabelina, D., Saporta, A., & Beeman, M. Flexible or leaky attention in creative people? Distinct patterns of attention for different types of creative thinking. Memory & cognition, 44(3), 488–498 (2016).

66. Boot, N., Baas, M., Mühlfeld, E., de Dreu, C. K., & van Gaal, S. (2017). Widespread neural oscillations in the delta band dissociate rule convergence from rule divergence during creative idea generation. Neuropsychologia, 104, 8–17 (2017).

67. Jann, K., Kottlow, M., Dierks, T., Boesch, C., & Koenig, T. Topographic electrophysiological signatures of fMRI resting state networks. PloS one, 5(9), e12945. (2010).

68. Guilford, J. P. Creativity: Yesterday, today and tomorrow. The Journal of Creative Behavior, 1(1), 3–14 (1967).

69. Mednick, S. The associative basis of the creative process. Psychological review, 69(3), 220 (1962).

70. Torrance, E. P. (1974). The Torrance Tests of Creative Thinking-Norms Technical Manual Research Edition-Verbal Tests, Forms A and B Figural Tests, Forms A and B. Princeton, NJ: Personnel Press.

71. Immordino-Yang, M. H., Christodoulou, J. A., & Singh, V. Rest is not idleness: Implications of the brain’s default mode for human development and education. Perspectives on Psychological Science, 7(4), 352–364 (2012).

72. Landis, J. R., & Koch, G. G. An application of hierarchical kappa-type statistics in the assessment of majority agreement among multiple observers. Biometrics, 363–374 (1977).

73. Vigário, R. N. (1997). Extraction of ocular artefacts from EEG using independent component analysis. Electroencephalography and Clinical Neurophysiology, 103, 395–404 (1997).

74. Kalcher, J., Pfurtscheller, G. Discrimination between phase-locked and non-phase locked event-related EEG activity. Electroencephalography and clinical neurophysiology, 94, 381–384 (1995).

75. Kloosterman, N. A., Meindertsma, T., Hillebrand, A., van Dijk, B. W., Lamme, V. A., & Donner, T. H. Top-down modulation in human visual cortex predicts the stability of a perceptual illusion. Journal of Neurophysiology, 113, 1063–1076. (2015).

76. Donner, T. H., Siegel, M. A framework for local cortical oscillation patterns. Trends in Cognitive Sciences, 15, 191–199 (2011).

77. Mitra, P. P., & Pesaran, B. Analysis of dynamic brain imaging data. Biophysical journal, 76(2), 691–708 (1999).

78. Oostenveld, R., Fries, P., Maris, E., Schoffelen, J. M. FieldTrip: open source software for advanced analysis of MEG, EEG, and invasive electrophysiological data. Computational Intelligence and Neuroscience, 2011, 156869 (2011).

79. Monin, Benoît, and Daniel M. Oppenheimer. “Correlated averages vs. averaged correlations: Demonstrating the warm glow heuristic beyond aggregation.” Social Cognition 23.3, 257–278 (2005).

80. Benjamini, Y., & Hochberg, Y. Controlling the False Discovery Rate: A Practical and Powerful Approach to Multiple Testing. Journal of the Royal Statistical Society, 57(1), 289–300 (1995).

